# Ecological context gates numerosity-based affiliation decision in zebrafish

**DOI:** 10.64898/2026.02.08.704638

**Authors:** Hsin-En Cheng, Christoph D. Dahl

## Abstract

In many group-living animals, affiliating with larger conspecific aggregations can reduce predation risk through dilution. A central question is what simple mechanisms can support such affiliation: are responses driven by generic visual magnitude cues (e.g., “more visual mass” results in stronger attraction), or is affiliation conditioned on perceptual cues about what is being viewed, such that numerical group-size information is used for conspecific-like stimuli but suppressed for predator-like stimuli. Here we test this in zebrafish (*Danio rerio*) by orthogonally manipulating number and stimulus regime (conspecific zebrafish line drawings versus predator-like large-fish line drawings, differing in both apparent body size and body morphology/identity). In a spontaneous-choice assay with high-resolution tracking, zebrafish preferentially affiliated with the larger of two groups when stimuli were conspecific-like, and preference strength scaled with the logarithmic ratio between groups, consistent with ratio-sensitive numerical processing of group size. Trial-level model comparison favoured an Approximate Number System (ANS) predictor over an object-tracking (OTS) “small-number” account, with no additional small-number advantage under these conditions. When stimuli were predator-like, numerosity-based affiliation was abolished and sometimes weakly reversed, and neither ANS nor OTS predictors explained systematic variance, consistent with threat-like appearance suppressing affiliation even when a larger group is available. These findings show that zebrafish display ANS-like sensitivity to conspecific group size, but that its behavioural influence is selectively deployed: a predator-like stimulus regime (larger body size with correlated changes in morphology/identity) gates whether numerical information guides social affiliation.

## Introduction

Discriminating between quantities is a core cognitive ability in animals, with particular relevance to social species in which group size impacts predation risk and resource access. Across taxa, animals show consistent and ecologically meaningful sensitivity to number. Rhesus macaques treat *number* as an abstract organizing dimension, spontaneously matching displays by numerosity over competing visual features (e.g., color, shape, surface area), with performance improving at easier numerical ratios (Cantlon and Brannon, 2007). Capuchin monkeys reliably discriminate larger quantities across both food and token stimuli, with performance scaling by numerical ratio, indicating use of an analogue magnitude system independent of stimulus type (Addessi et al., 2008). Prosimian primates also spontaneously select the larger of two food quantities when ratios are sufficiently large (e.g., 1 vs 3), doing so across both small and large set sizes (Jones and Brannon, 2012). This suggests that ratio-based numerical discrimination predates the divergence of higher primates. In dogs, visual studies consistently find spontaneous choice of the larger visible amount with ratio-dependent performance (Ward and Smuts, 2007; Petrazzini and Wynne, 2016), whereas olfactory tests show robust detection but only limited quantity discrimination in untrained animals (Horowitz et al., 2013), with performance modulated by cephalic index (i.e., skull shape) (Ferrando and Dahl, 2022). Newborn chicks map smaller numerosities to the left and larger to the right relative to a reference set, indicating an early, spatially organized representation of number (Rugani et al., 2015). Even insects display numerical competence: honeybees spontaneously discriminate quantity, preferring the larger set when the smaller set is one and the numerical distance is sufficiently large (Howard et al., 2020). Together, these findings indicate that approximate quantity representations are widespread and functionally relevant across diverse taxa. To investigate how such representations are deployed in a specific ecological context, we focus on teleost fish.

In teleost fish, group association provides a natural test case: larger groups offer reduced individual predation risk via dilution and confusion effects, while also improving access to resources (Krause and Ruxton, 2002). Numerous studies show that fish such as zebrafish, guppies, and cichlids can distinguish between groups of different sizes, often exhibiting a preference for larger groups (Agrillo et al., 2008; Gómez-Laplaza and Gerlai, 2011; Agrillo et al., 2010; Bisazza et al., 2010; Agrillo et al., 2012). This preference is typically ratio-dependent and follows Weber-like scaling (Gómez-Laplaza and Gerlai, 2011; Agrillo et al., 2010). Early studies often relied on training and conditioning, associating group size with rewards. While these demonstrate the capacity for quantity discrimination, reinforcement paradigms risk conflating learned strategies with spontaneous behaviour (Agrillo et al., 2008; Gómez-Laplaza and Gerlai, 2011). Accordingly, researchers have increasingly adopted spontaneous-choice tests, in which naïve fish choose freely between two group options without reinforcement. These designs reveal untrained tendencies and likewise show a consistent preference for larger groups, with stronger effects as numerical disparity increases (Agrillo et al., 2008; Dadda et al., 2009; Agrillo et al., 2017; Agrillo and Bisazza, 2018). At the same time, considerable debate remains over whether observed choices reflect true numerical processing or reliance on non-numerical cues such as cumulative area, density, convex-hull area, or motion (Gebuis and Reynvoet, 2011; Nys and Content, 2012). While some studies attempt to equate these cues across conditions, others accept that numerosity is often embedded within multiple correlated dimensions. However, numerical information does not necessarily dominate group-association choices: zebrafish integrate group size with other social cues, such as familiarity, and can shift from preferring familiar groups to larger unfamiliar groups once size differences become sufficiently large (Swaney et al., 2025). More generally, cues associated with predation threat can suppress group approach and thereby attenuate or abolish numerosity-based preferences.

A notable limitation of most previous work is that stimuli are conspecific-like: live groups, video or animated conspecifics, or dot arrays designed to mimic small fish. Far less is known about how predator-like visual regimes, in which apparent size covaries with morphology/identity, modulate group-association choice. Yet body size is ecologically meaningful: large individuals can signal threat, dominance, or predation risk. In zebrafish, looming or large angular size recruits escape circuits (Dunn et al., 2016), and even computer-animated images of sympatric predators such as the Indian leaf fish elicit robust fear-like responses (Gerlai et al., 2009; Luca and Gerlai, 2012). Classic studies of oddity and size-assortative group association further show that fish avoid unusually large heterospecifics in mixed groups (Lima and Dill, 1990; Landeau and Terborgh, 1986; Peuhkuri, 1997). To our knowledge, no study has tested how such size cues interact with numerosity preferences. Thus, if large fish stimuli signal danger, affiliation should be attenuated or reversed, consistent with predation-risk theory and risk-sensitive group association (Lima and Dill, 1990; Kelley and Magurran, 2003; Krause and Ruxton, 2002). From a predation-risk perspective, decision rules should trade off affiliation benefits against perceived threat, so stimulus meaning (e.g., apparent body size) is expected to gate approach-avoidance (Lima and Dill, 1990). Consistent with this view, group association choices in fishes are modulated by risk cues and perceived safety (Kelley and Magurran, 2003; Krause and Ruxton, 2002; Mukherjee and Bhat, 2023).

Here we address this gap by asking not only whether zebrafish represent relative group size, but when those representations guide overt decisions and how they are modulated by ecologically meaningful variation in apparent body size and associated predation risk. Using a spontaneous-choice paradigm with static visual stimuli, we manipulated both the number and apparent size of stimulus fish. Static image cues such as cumulative area and convex hull were quantified and included as covariates in the analyses. Thus, our findings reflect quantity preferences that are modulated by ecological context rather than being reducible to simple low-level magnitude cues.

Zebrafish are a particularly suitable model for studying quantitative decision-making. They display robust group-association behaviour and show larger-group preferences with clear ratio limits under spontaneous choice (Qin et al., 2014; Pritchard et al., 2001). Beyond discrimination, recent work reports proto-arithmetic operations in zebrafish, further supporting the suitability of small teleosts for mechanistic studies of quantitative cognition (Potrich et al., 2024). In addition, two-dimensional stimuli reliably evoke social affiliation, enabling tightly controlled visual presentations while retaining ecological validity (Saverino and Gerlai, 2008), consistent with spontaneous-choice evidence for ratio-dependent preferences in fishes (Agrillo et al., 2008; Dadda et al., 2009; Agrillo et al., 2017; Agrillo and Bisazza, 2018). Zebrafish are therefore widely used in both behavioural and neuroethological research on social cognition, making them an excellent system in which to investigate how numerosity-guided affiliation is modulated by ecologically meaningful cues.

We use the term *affiliation preference* to denote greater occupancy near the side depicting higher numerosity when stimuli are conspecific-like (small). By contrast, with predator-like (large) stimuli we refer to *risk-avoidance*, i.e., reduced occupancy near stimulus sides (and sometimes increased central occupancy). These definitions align with established distinctions between group attraction and predator avoidance responses in zebrafish.

At an ultimate level, affiliation with larger groups is expected because larger groups provide adaptive social and antipredator benefits. Thus, a straightforward prediction is that preference for the larger group should increase with numerical disparity (ratio dependence). At a proximate level, however, such behaviour could be mediated by continuous visual magnitudes that covary with number, such as cumulative area, density, convex-hull area, or perimeter. In this framework, cumulative area provides a clear prediction: groups with greater total surface area should elicit stronger affiliation, regardless of the apparent body size of the individuals. By contrast, if the apparent size of the fish carries ecological meaning (as implemented here via conspecific-like vs. predator-like fish line-drawing models), then affiliation preferences are expected to be modulated by stimulus size: conspecific-like groups should attract, whereas predator-like groups may attenuate or even reverse affiliation, consistent with risk-avoidance. In addition, we include one boundary condition: in equal-number contrasts there is no objectively “correct” side, so the key question is whether fish affiliate with one group rather than remain centrally.

As most numerosity studies have used conspecific-like stimuli (live/animated shoals or dot arrays) (Agrillo and Bisazza, 2018; Agrillo et al., 2017), our design explicitly pushes apparent body size into predator-like ranges while manipulating number. When body size is manipulated, it is typically to probe social preference or spacing rather than numerosity per se (e.g., number-by-size manipulations in animations, preferences for larger conspecifics, or systematic reviews of social tests and spacing) (Fernandes et al., 2015; Aslanzadeh et al., 2019; Ogi et al., 2021; Pita and Fernández-Juricic, 2021). To our knowledge, no studies have pushed apparent body size into *predator-like* ranges while simultaneously testing numerosity choices (Agrillo and Bisazza, 2018; Agrillo et al., 2017). Neuroethological work shows that looming or large angular size recruits escape circuits in zebrafish (Temizer et al., 2015; Dunn et al., 2016; Fotowat and Engert, 2023), and classic predation studies document oddity and size-assortative group association (Landeau and Terborgh, 1986; Peuhkuri, 1997). Together, these findings imply that apparent size can fundamentally reverse approach tendencies.

A further question is *how* number is represented. Across species, two complementary regimes are often discussed (Feigenson et al., 2004; Dehaene, 2011; Gallistel and Gelman, 2000): an *Approximate Number System* (ANS) that yields Weber-like, ratio-dependent discrimination, and a small-number *object-tracking* regime (OTS) that can confer an advantage for very small sets (typically ⩽ 4). In fish, spontaneous-choice and training studies broadly support ANS-like ratio effects at larger numerosities (e.g., Agrillo et al., 2010; Gómez-Laplaza and Gerlai, 2011; Agrillo et al., 2012; Agrillo and Bisazza, 2018), with mixed evidence for an additional boost in the subitizing range, especially when non-numerical cues are tightly controlled (Agrillo and Bisazza, 2018; Gebuis and Reynvoet, 2012). Recent work in the miniature teleost *Danionella cerebrum* extends this picture: using a habituation–dishabituation paradigm with dot arrays in which basic continuous visual properties were systematically controlled (Zanon et al., 2025). Zanon et al. (2025) found robust dishabituation when numerosity changed (e.g. 3 vs. 9 elements), reinforcing that fish can extract numerical information even when simple magnitude cues are constrained. Our size manipulation provides a principled test of these regimes in an ecologically grounded context: if zebrafish recruit an OTS-like process, any small-number advantage should appear with conspecific-like stimuli but *not* with predator-like stimuli, where risk-avoidance is expected to suppress numerosity-based affiliation. It is important to note that recent syntheses emphasise that behavioural signatures often reflect mixtures of representational and decision processes, so ANS/OTS should be treated as useful models rather than mutually exclusive mechanisms (Nieder, 2025). Accordingly, we use ‘ANS/OTS’ as behavioural model classes rather than as direct claims about neural representational format.

We examine spontaneous quantity discrimination while manipulating both numerosity and stimulus regime (conspecific zebrafish line drawings vs. predator-like large-fish line drawings). Regimes differ in apparent body size and fish-model morphology/identity). Specifically, we test four primary predictions: (i) For conspecific-like stimuli, *affiliation preference* should increase with the log-ratio of numerosity, reflecting a larger-group advantage (H1). (ii) For predator-like stimuli, we predict an *ecological-gating* effect: affiliation should deviate from this pattern, being attenuated, absent, or reversed, consistent with risk-avoidance (H2). Such divergence would indicate that continuous magnitude cues cannot account for behaviour across the full stimulus set, since those magnitudes increase with numerosity regardless of size. Conversely, if larger groups are still preferred when predator-like, the data would instead be parsimoniously explained by continuous visual magnitudes. (iii) In equal-number contrasts, we predicted that fish would show lateral preference and group association behaviour with small stimuli, but remain more centrally positioned with large stimuli (H3). In addition, we formulate a model-based hypothesis that links these effects to ANS/OTS accounts: (iv) In unequal-number trials, preference strength should scale with numerical *ratio* (ANS). Any extra small-number improvement, if OTS applies, should be restricted to pairs composed only of very small sets (1, 2, or 4) and should be observable for conspecific-like stimuli but not for predator-like stimuli (H4). As a manipulation check, we include single-option contrasts (0 vs. N) to assess whether conspecific-like groups elicit social approach and whether this tendency is abolished or reversed for predator-like stimuli.

Because continuous visual magnitudes can covary with number (e.g., cumulative area, density, convex hull), a magnitude-based account predicts that association should vary monotonically with these metrics and generalise across stimulus size. By contrast, an ecologically grounded account predicts that predator-like stimulus appearance (including apparent size and morphology) modulates valuation, yielding a Size×Disparity interaction: the same numerical (or area) disparity produces stronger affiliation for conspecific-like groups but attenuated or reversed affiliation for predator-like stimuli.

## Materials and methods

### Subjects

Experiments were conducted on a total of 30 adult zebrafish (*Danio rerio*, AB strain). Eighteen fish performed the single-option (0 vs *N* ; see details below) and twelve fish performed the two-choice (number-vs-number) cohort, which included both equal and unequal contrasts (i.e., *N*_L_ vs. *N*_R_). The fish were housed in 35 × 22 ×25 cm (L × W × H) glass aquariums with continuous water exchange (dechlorinated tap water, 5% daily) at 28.5°C and subjected to the standard 14h/10h cycle. Subjects’ ages ranged from 12 to 18 months. The fish were held at the core laboratory of zebrafish (CLZ) at the Department of Anatomy and Cell Biology of Taipei Medical University, Taiwan. All procedures complied with institutional and national ethical guidelines.

### Experimental apparatus

Testing was conducted in a custom-built rectangular aquarium (40 × 15 × 18 cm; L × W × H), filled with dechlorinated water maintained at 28.5°C to a depth of 17 cm. The aquarium was divided into three compartments: a central neutral zone (L: 20 cm) and two side compartments (L: 10 cm), separable via acrylic sliders. Each side compartment was equipped with a slide-in mount on the outer wall, allowing for the insertion of laminated, printed stimulus displays. To minimize external distractions and control lighting conditions, the aquarium was placed inside an enclosure (60 × 60 × 90 cm; L × W × H) constructed from white polypropylene panels mounted on a T-slotted aluminium frame (20 Series; Misumi Group Inc., Tokyo, Japan). The enclosure featured a door that was opened between trials for access and closed during each trial to ensure constant illumination and prevent outside visual cues. No environmental enrichment was provided during testing. The long sides of the aquarium were rendered opaque using white acrylic panels to block extraneous visual input.

### Stimuli

Each stimulus depicted a static group of black line drawings of fish presented on a neutral white background. Stimuli represented groups of 1, 2, 4, 8, or 16 individuals viewed from a lateral perspective. Two stimulus regimes were used. In the *conspecific-like* (small) regime, each individual was a line drawing of an adult zebrafish (length ≈ 25 mm, width ≈ 6 mm). In the *predator-like* (large) regime, each individual was a line drawing parameterised to approximate a likely zebrafish predator, based on a juvenile snakehead (*Channa* sp.; length ≈ 47 mm, width ≈ 18 mm) (Lima and Dill, 1990; Kelley and Magurran, 2003). We did not include an independent manipulation check to confirm predator categorisation in the present setup. Across stimulus regimes, numerosity and the spatial arrangement of individuals within each display were held constant: for each numerosity, the same layout was used, and only the fish model changed between regimes (conspecific vs. predator line drawing, with corresponding differences in apparent body size). Because the two regimes differ in fish-model morphology/identity as well as apparent size, we treat this manipulation as an ecologically motivated conspecific-like vs. predator-like stimulus regime, not as body size in isolation. This design holds numerosity and layout fixed while varying stimulus appearance across regimes, allowing us to test whether predator-like large-fish displays modulate choice independently of numerosity and spatial arrangement. Stimuli were printed in grayscale at high resolution, laminated for waterproofing, and inserted into custom 3D-printed holders mounted externally along the long sides of the test tank. Example stimuli for both regimes are shown in Fig. 1A,D.

**Figure 1:**
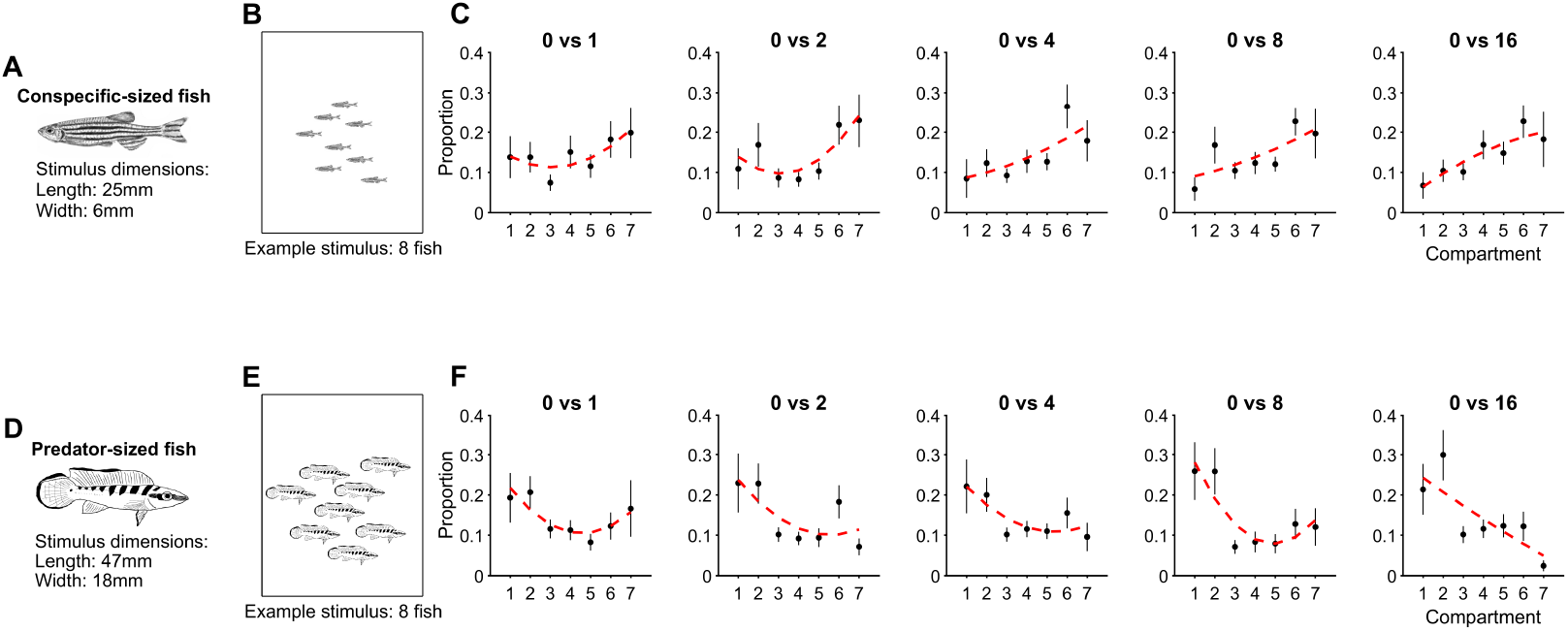
Compartment occupancy profiles in single group presence. (A) Line drawing of the conspecific-like stimulus fish (dimensions correspond to adult zebrafish body size). (B) Example static display of a group of eight conspecific-like stimulus fish. (C) Spatial occupancy profiles when only one side of the tank contained a group of small conspecific-like fish (0 vs. 1, 0 vs. 2, 0 vs. 4, 0 vs. 8, 0 vs. 16; top row). (D) Line drawing of the predator-like stimulus fish. (E) Example display of eight predator-like stimulus fish. (F) Spatial occupancy profiles for zero-versus-group trials with predator-like fish (0 vs. 1, 0 vs. 2, 0 vs. 4, 0 vs. 8, 0 vs. 16; bottom row). In (C) and (F), dots show the mean proportion of time spent in each of the seven virtual compartments; error bars indicate SEM. Red dashed curves show the best-fitting polynomial (linear or quadratic) to the compartment means.

### Stimulus feature extraction (image-based controls)

Fish line drawings were segmented by selecting the cleaner of a global vs. adaptive threshold (edge-score criterion), followed by light morphology (opening/closing, hole fill) and a robust “small→big” merge that dilates large components and absorbs nearby fragments before relabeling. From the final connected components we computed: object count (*N*), coverage, and mean nearest-neighbour spacing. Coverage was defined as the fraction of image pixels occupied by fish, 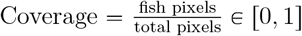. Nearest-neighbour spacing was computed from component centroids 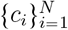 as 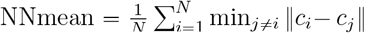 (in pixels), defined only when *N* ⩾ 2 (see GLMM for handling of missing values). Metrics were computed separately for the *small* and *large* stimulus sets and summarised by numerosity and side (L/R). For each behavioural contrast with numerosities *N*_1_ (Left) and *N*_2_ (Right), we derived covariates. Note that the Left/Right alignment follows the relative magnitude within each contrast (i.e., *N*_1_ < *N*_2_), not the actual stimulus position; stimulus sides were counterbalanced across trials.

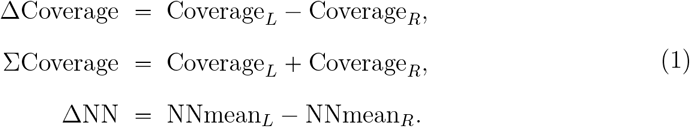

When a side-specific spacing value was unavailable (e.g., *N* <2), ΔNN was treated as missing at the modelling stage (mean imputation with a missingness indicator; see Eq. (8)). These image-based covariates were included only as controls in the statistical models.

### Experimental protocol

At the beginning of each trial, a single test fish was gently placed in the central (neutral) compartment of the aquarium and allowed to acclimate (for ≈ 5 mins). Once acclimated, laminated stimulus sheets (representing the designated group size) were inserted into the slide-in mounts on the exterior sides of the two side compartments. The acrylic sliders separating the compartments were then carefully removed, allowing the fish free access to the entire tank for a 1-minute observation period. During this time, the fish’s movements and compartment choices were continuously recorded.

At the end of the trial, the acrylic sliders were reinserted to confine the fish to the central compartment, and the stimulus sheets were removed from the side compartments. The tank was then prepared for the next trial, either with a new set of stimuli or a new subject as required by the experimental design. Two independent cohorts were used: a two-choice cohort (number-vs-number; *N*_L_ vs. *N*_R_, including equal and unequal contrasts) and a zero–vs–*N* cohort (18 subjects). For two–choice tests, each of 12 subjects completed all 15 pairwise contrasts for each stimulus size, yielding *n* = 180 trials per size (small and large). These comprise 120 unequal–number trials (10 contrasts/subject) and 60 equal–number trials (5 contrasts/subject). For zero–vs–*N* tests, each of 18 subjects completed the five *N* levels per size, yielding *n* = 90 trials per size. All scoring and tracking were automated.

### Data acquisition and analysis

Fish movements were recorded using a FLIR^®^ Grasshopper USB3 camera (GS3-U3-28S4C; 1/1.8” Sony ICX687 sensor; 1928 × 1448 pixel resolution) equipped with a 12 mm C Series lens (Edmund Optics^®^, Stock #58-001), mounted perpendicular to the water surface at a distance of 80 cm above the aquarium. Uniform illumination inside the enclosure was provided by a Mettle^®^ SL400 LED panel (45 W, 2100 lm, 350 × 250 mm), positioned 25 cm above the top panel of the enclosure. Camera control and image acquisition were performed with MATLAB (Mathworks^®^, Natick, MA, USA) using the Image Acquisition and Image Processing Toolboxes. The lens aperture was set to f/4.5 to maximize depth of field and image sharpness. Video recordings were acquired at 26 frames per second using custom-written scripts in MATLAB and the Image Acquisition Toolbox.

Fish trajectories in the x-y plane were extracted from the recorded videos and post-processed according to a custom pipeline. In a first step, a background subtraction and object detection was applied: For each video frame, a static background model was computed, and each frame was subtracted from this background to identify moving objects. Thresholding and morphological filtering were applied to isolate the fish as a single contiguous blob. The centroid of the detected fish blob was calculated for each frame, yielding a time series of x-y coordinates that represented the fish’s position throughout the trial. The x-y coordinates were then mapped to one of seven equally spaced compartments (bins) along the tank’s long axis. For each trial, the trajectory (x-y coordinates), per-frame compartment assignment, and total occupancy per compartment were saved for subsequent analysis. The position of the fish was tracked frame by frame throughout the 1-minute observation period, allowing for quantification of spatial preferences, compartment occupancy, and laterality indices. All steps were performed automatically in batch mode, without manual intervention or subjective corrections.

All statistical analyses were performed using custom-written scripts in MATLAB. The primary aim of the analysis was to quantify spatial preferences and side choices of test fish in response to different numerical contrasts presented in the two side compartments. Trials were categorized according to the number of stimulus fish presented on the left and right sides, as well as the size of the stimulus fish (*small* vs *large*). For each condition (e.g., 1 vs 4 small fish), the occupancy per compartment was computed by averaging the proportion of time spent in each compartment across all trials (equal to the number of subjects) belonging to the same condition. The standard error of the mean (SEM) was calculated for each bin to provide an estimate of variability. For trials in which the numerical contrast was equal on both sides (e.g., 8 vs. 8), a correction procedure was applied to account for the fact that there is no correct or expected side preference in these conditions. From the perspective of the focal fish, either side represents an equally valid choice, as both offer the same numerical stimulus. The correction consisted of flipping individual compartment profiles as necessary so that, across all trials, the direction of preference (toward either side) is consistently represented. This approach allows us to measure the strength of a fish’s decision to approach a group, regardless of which side was chosen. Choice alignment (mirroring) was used only for descriptive summaries of equal-number contrasts (*N*_left_ = *N*_right_). All inferential analyses were performed on the original trial data. When a signed ratio predictor or an accuracy label was required, unequal-number contrasts were magnitude-aligned (swapping left/right labels so that the larger numerosity was assigned to the right), but no choice alignment was applied. Equal-number contrasts were omitted only from analyses that require a well-defined larger side (e.g., the accuracy GLMM and the ANS/OTS model comparison restricted to ratio > 1). From each compartment profile, two shape descriptors were extracted by polynomial fitting. A linear fit yielded the *slope*, reflecting overall directional bias across the tank axis, while a quadratic fit yielded the *curvature*, reflecting central versus peripheral attraction. Both coefficients, together with their *R*^2^ values, were retained for each condition. In parallel, for unequal-number two-choice trials we applied *magnitude-alignment* (swapping sides as needed so that the higher numerosity was on the right) and then computed a trial-level laterality index (LI) as the difference in mean occupancy between the two bins adjacent to the higher-numerosity group and the corresponding two bins adjacent to the lower-numerosity group:

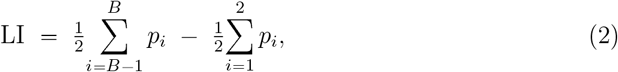

where *p*_*i*_ is the proportion of time spent in bin *i* and bins are indexed left → right, with *B* = 7 the rightmost bin. Positive values indicate preference for the larger group, negative values indicate preference for the smaller group, and values near zero suggest no consistent choice.

Group-level results are reported as mean ± SEM across trials. Relationships between slope, curvature, and laterality index and the numerical ratio of groups were examined using linear regression. Ratios were log_2_-transformed to account for Weber-like scaling of discrimination performance. Statistical comparisons of laterality indices against zero were performed with Wilcoxon signed-rank tests. For single-option (0 vs. *N*) contrasts, the primary one-sided hypotheses were LI > 0 for conspecific-like (small) stimuli and LI < 0 for predator-like (large) stimuli. For completeness we also report the opposite-tail and two-sided *p*-values. Per-*N* tests were Bonferroni-corrected within each stimulus-size class across the five tested numerosities (*α* = 0.05/5 = 0.010).

Results were visualized as mean ± SEM profiles across compartments for each condition, as well as scatterplots showing laterality indices and regression lines to illustrate the dependence of group preference on the numerical ratio. Discrimination performance in numerosity tasks is generally ratio-dependent rather than difference-dependent, following Weber’s law (Feigenson et al., 2004; Agrillo and Bisazza, 2018; Messina et al., 2021). That is, animals tend to discriminate two sets more easily when the *relative difference* between them is large (e.g., 2:4 = ratio 0.5) than when it is small (e.g., 8:10 = ratio 0.8), even though both contrasts differ by two items in absolute terms. Formally, Weber’s law implies that the discriminability of two numerosities *n*_1_ and *n*_2_ depends on their ratio rather than their absolute difference. This can be expressed as:

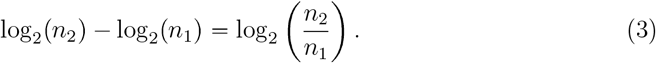

Accordingly, we analysed numerical contrasts in terms of their log_2_ ratio, which linearises relative differences and provides an intuitive scale: for example, 2 vs. 4 (ratio 2: 1) and 4 vs. 8 (ratio 2: 1) both yield log_2_ ratio = 1, predicting equal discriminability under Weber-like scaling. In practice, many species, including fish, sometimes show disproportionately strong performance at very small numerosities (e.g., 1 vs. 2, 2 vs. 4) (Agrillo et al., 2012; Piffer et al., 2012). Such deviations from Weber’s law are often interpreted as evidence for an additional object-tracking system (OTS), which provides precise representations of small sets, operating alongside the approximate number system (ANS) that supports ratio-dependent estimation at larger set sizes. To evaluate whether zebrafish quantity discrimination was driven solely by Weber-like scaling or showed additional advantages at small numerosities, we fitted competing regression models. The dependent variable was the laterality index (LI) computed at the trial level, quantifying preference for the larger versus smaller group. Two predictors were considered. First, an *ANS predictor* was defined as the log_2_ ratio of the two numerosities:

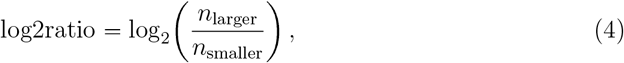

which captures Weber-like dependence on relative rather than absolute differences. Second, an *OTS predictor* was defined as a binary factor coding whether both numerosities fell within the small range (1, 2, or 4) versus included larger values (> 4):

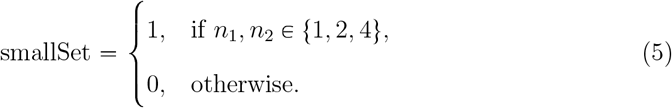

This predictor tests whether discrimination at small set sizes was disproportionately strong, consistent with object-tracking mechanisms. For the OTS–ANS model comparison, equal-number contrasts (ratio = 1) were excluded, and log_2_ ratios were *z*-scored to improve numerical conditioning. We fitted (i) an ANS model (LI ∼ log_2_ ratio), (ii) an OTS model (LI ∼ smallSet, with smallSet = 1 if both numerosities ∈ { 1, 2, 4 }), and (iii) a combined model (LI ∼ log_2_ ratio + smallSet). Model fit was evaluated via Akaike Information Criterion (AIC) and adjusted *R*^2^. The contribution of smallSet in the combined model was assessed by the coefficient *p*-value.

### Linear mixed-effects model (trial-level laterality)

To test whether laterality increased systematically with group-size ratio, we fitted linear mixed-effects models (LMEs) to the per-trial laterality index (LI) for both conspecific-like (*small*) and predator-like (*large*) stimuli:

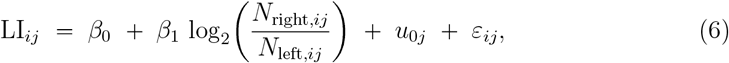

where *u*_0*j*_ is a random intercept for subject *j* (12 individuals) and *ε*_*ij*_ is the residual. For unequal-number trials, records were magnitude-aligned so that the larger numerosity was assigned to the right (sides swapped if necessary) before computing log_2_(*N*_right_/*N*_left_). For equal-number trials (*N*_right_ = *N*_left_), no magnitude alignment was applied and these trials contribute to the model at log_2_(*N*_right_/*N*_left_) 0. Models were fitted by maximum likelihood in MATLAB (`fitlme`). We report fixed-effect estimates (estimate ± SE), 95% confidence intervals, *p*-values, and marginal/conditional *R*^2^ following Nakagawa and Schielzeth (2013). Predicted population means for the LI–vs–log_2_ ratio panels were obtained at ratios 0–4 using the function `predict` with ‘ `Conditional’, false`. Here and throughout, “right” is a *labeling convention* denoting the side carrying the larger numerosity after normalization.

### Size × ratio interaction model

To test whether ratio-dependence differed between conspecific-like (small) and predator-like (large) stimuli, we fitted a pooled linear mixed-effects model including size, log_2_ ratio, and their interaction, with subject-specific random intercepts and random slopes for log_2_ ratio:

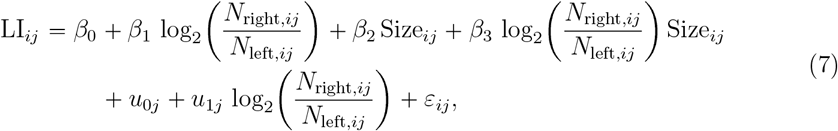

where Size_*ij*_ indicates stimulus regime (small/conspecific-like line drawings as the reference level). Because log_2_(*N*_right_/*N*_left_) is undefined when one side is zero, the interaction analysis was restricted to two-choice trials with *N*_left_ > 0 and *N*_right_ > 0. Equal-number contrasts were retained and contribute at log_2_(*N*_right_/*N*_left_) = 0 (no magnitude alignment is applicable when *N*_left_ = *N*_right_). Models were fitted by maximum likelihood in MATLAB (`fitlme`). Evidence for the interaction was evaluated both via the fixed-effect interaction term and via a likelihood-ratio comparison to the corresponding main-effects model.

In a further step, we tested whether numerical laterality persists after controlling for low-level properties of the printed stimuli at two complementary levels.

#### (i) Trial-level GLMM on choice accuracy

For unequal-number *two-choice* trials (*N*_left_ > 0, *N*_right_ > 0, and *N*_left_ ≠ *N*_right_), we first applied a *magnitude-alignment* step by swapping left/right labels as needed so that the larger numerosity was assigned to the right. We then coded Accuracy = 1 if the fish chose the side containing the larger numerosity and 0 otherwise, and used the aligned log-ratio predictor log_2_(*N*_right_/*N*_left_) (and its *z*-scored version) in the model. Equal-number trials (*N*_left_ = *N*_right_) have no objectively larger side and therefore no well-defined accuracy label. Accordingly, they were not included in this accuracy GLMM. These equal-number trials were analysed separately on the original (raw; not choice-aligned) data, and we additionally report a *choice-aligned* descriptive summary to quantify the strength of side commitment irrespective of whether the left or right side was chosen. We fit a binomial GLMM (logit link, Laplace) as:

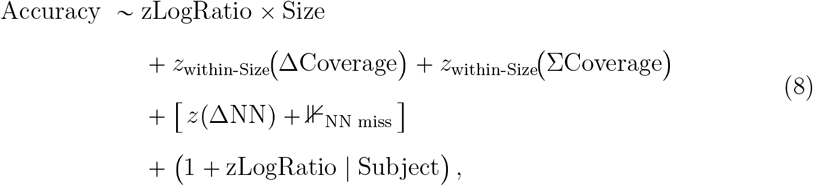

where zLogRatio = *z* log_2_(*N*_2_/*N*_1_) and *Size* indicates stimulus regime (*small* /conspecific-like line drawings vs. *large*/predator-like line drawings). Coverage covariates were centred and scaled *within* size class. When available, ΔNN was *z*-scored with mean imputation and a missingness indicator. Fixed-effect coefficients are reported with SEs, *p*-values, and odds ratios. Because numerosity and coverage were naturally correlated in our displays, this specification is deliberately conservative: zLogRatio competes with highly collinear coverage terms, trading off sensitivity to numerical effects against stringent control for simple continuous magnitudes. We therefore interpret this GLMM primarily as a check that no single coverage or spacing metric can trivially account for choice behaviour, rather than as the most sensitive estimate of numerosity effects.

#### (ii) Condition-level regression on laterality

Independently, trials were aggregated by condition and size class, and the condition mean laterality index (LI; larger-side minus smaller-side edge occupancy) was regressed on numerical ratio and image covariates (defined in Eq. (1)):

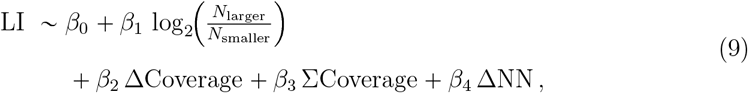

fitting *small* and *large* stimulus sets *separately*. This tests Weber-like, ratio-dependent laterality while adjusting for coverage/spacing asymmetries. Coefficients are reported as estimate ± SE with two-sided *p*-values.

Together, these analyses ensure that any numerical preference is not reducible to simple differences in occupied area or element spacing between sides.

## Results

### Single-option tests

We first assessed preference in zebrafish for social groups in the most basic context, i.e. trials in which only one side of the test tank contained an image of fish while the opposite side was left blank. When presented with groups of small, conspecific-like stimulus fish (Fig. 1A,B), zebrafish spent more time in the compartment near the occupied side, reflected in a positive laterality index (LI) (Table 1, Fig. 1C). Median LI values were positive for all tested group sizes (*N* = 1, 2, 4, 8, 16), but none of the per-*N* tests reached the Bonferroni-corrected threshold (*α* = 0.010; Table 1). When data across all group sizes were pooled, the population-level tendency to associate with conspecific-like groups was robust (median = 0.106, *n* = 90, primary one-sided test LI > 0: *p* = 0.001828; two-sided *p* = 0.003656; Table 1). In contrast, when the same paradigm was used with groups of large, predator-like stimulus fish (Fig. 1D,E), zebrafish did not show approach toward the occupied side (Fig. 1F); laterality was negative across group sizes. Pooled across all group sizes, laterality was reliably below zero (median = −0.158, *n* = 90, primary one-sided test LI < 0: *p* = 0.000431; two-sided *p* = 0.000863; Table 1). At the per-*N* level, only *N* = 16 remained significant after Bonferroni correction (*p* = 0.00652), whereas the other group sizes did not (Table 1). This finding suggests a context-sensitive switch from *affiliation preference* for conspecific-like (small) stimuli to *risk avoidance* or neutrality for predator-like (large) stimuli (Table 1). To further characterise spatial occupancy with *conspecific-like (small)* stimuli, we fit linear and quadratic models to the mean compartment-occupancy profile for each contrast (Table 2, Fig. 1C). The linear slope parameter tended to increase with group size, indicating a gradual strengthening of affiliation preference as the number of stimulus fish increased, while quadratic curvature terms were small, consistent with a largely monotonic profile rather than pronounced peaking or centre avoidance. *R*^2^ values confirmed that the fits captured much of the systematic variance. To characterise spatial occupancy in trials with predator-like stimulus fish, we fit linear and quadratic models to the mean compartment-occupancy profiles for each 0 vs *N* contrast (Table 2, Fig. 1F). In these conditions, a consistently negative slope that becomes more negative with increasing group size, coupled with small positive curvature (*a* > 0), indicates an approximately monotonic decline in occupancy toward the stimulus that strengthens with group size, with only slight flattening near the stimulus (vertex outside the spatial range). This is consistent with risk-avoidance of predator-like groups.

**Table 1:** Zero vs. group: Laterality indices for small and large fish. Wilcoxon signed-rank tests of LI against zero with primary one-sided hypotheses (small: LI > 0; large: LI < 0). *p*-values are uncorrected; Bonferroni-corrected significance threshold for per-*N* tests within each size class is *α* = 0.010.

| $N$ | Small fish | | | Large fish | | |
| --- | --- | --- | --- | --- | --- | --- |
| | Median | $n$ | $p$ | Median | $n$ | $p$ |
| 1 | 0.021 | 18 | 0.2388 | -0.118 | 18 | 0.2567 |
| 2 | 0.184 | 18 | 0.1113 | -0.097 | 18 | 0.07534 |
| 4 | 0.138 | 18 | 0.0637 | -0.122 | 18 | 0.1113 |
| 8 | 0.123 | 18 | 0.0885 | -0.173 | 18 | 0.05353 |
| 16 | 0.085 | 18 | 0.0324 | -0.231 | 18 | 0.00652 |
| **All N** | **0.106** | **90** | **0.0018** | **-0.158** | **90** | **0.000431** |

**Table 2:** Summary of curve-fit parameters for 0 vs. *N* contrasts with small and large stimulus fish (raw means; bias-corrected means were identical to raw in this dataset). *Slope*: linear fit (*y* = *ax b*), coefficient *a. Curvature*: quadratic fit (*y* = *ax*^2^ *bx c*), coefficient *a. R*^2^: goodness-of-fit from the quadratic model.

| $N$ | Small fish | | | Large fish | | |
| --- | --- | --- | --- | --- | --- | --- |
| | Slope | Curvature | $R^2$ | Slope | Curvature | $R^2$ |
| 1 | 0.01115 | 0.00606 | 0.6457 | -0.01007 | 0.00893 | 0.7431 |
| 2 | 0.01703 | 0.00941 | 0.6688 | -0.02044 | 0.00654 | 0.5438 |
| 4 | 0.02141 | 0.00176 | 0.5682 | -0.01634 | 0.00581 | 0.6746 |
| 8 | 0.01953 | 0.00129 | 0.5417 | -0.02386 | 0.01323 | 0.7617 |
| 16 | 0.02271 | -0.00205 | 0.8132 | -0.03218 | 0.00062 | 0.6197 |

### Two-choice tests

We next tested preference when *both* sides of the tank contained groups of stimulus fish. With conspecific-like (*small*) stimuli, zebrafish generally spent more time near the larger group (Fig. 2B): the pooled laterality index (LI) across all pairwise contrasts was significantly greater than zero (median = 0.049, *n* = 180, *p* < 0.0001, Fig. 2C). By contrast, with predator-like (*large*) stimuli, ratio-dependent preference disappeared (Fig. 3B,C): LIs clustered near zero (sometimes slightly negative), consistent with reduced affiliation or mild avoidance under threat-like appearance.

**Figure 2:**
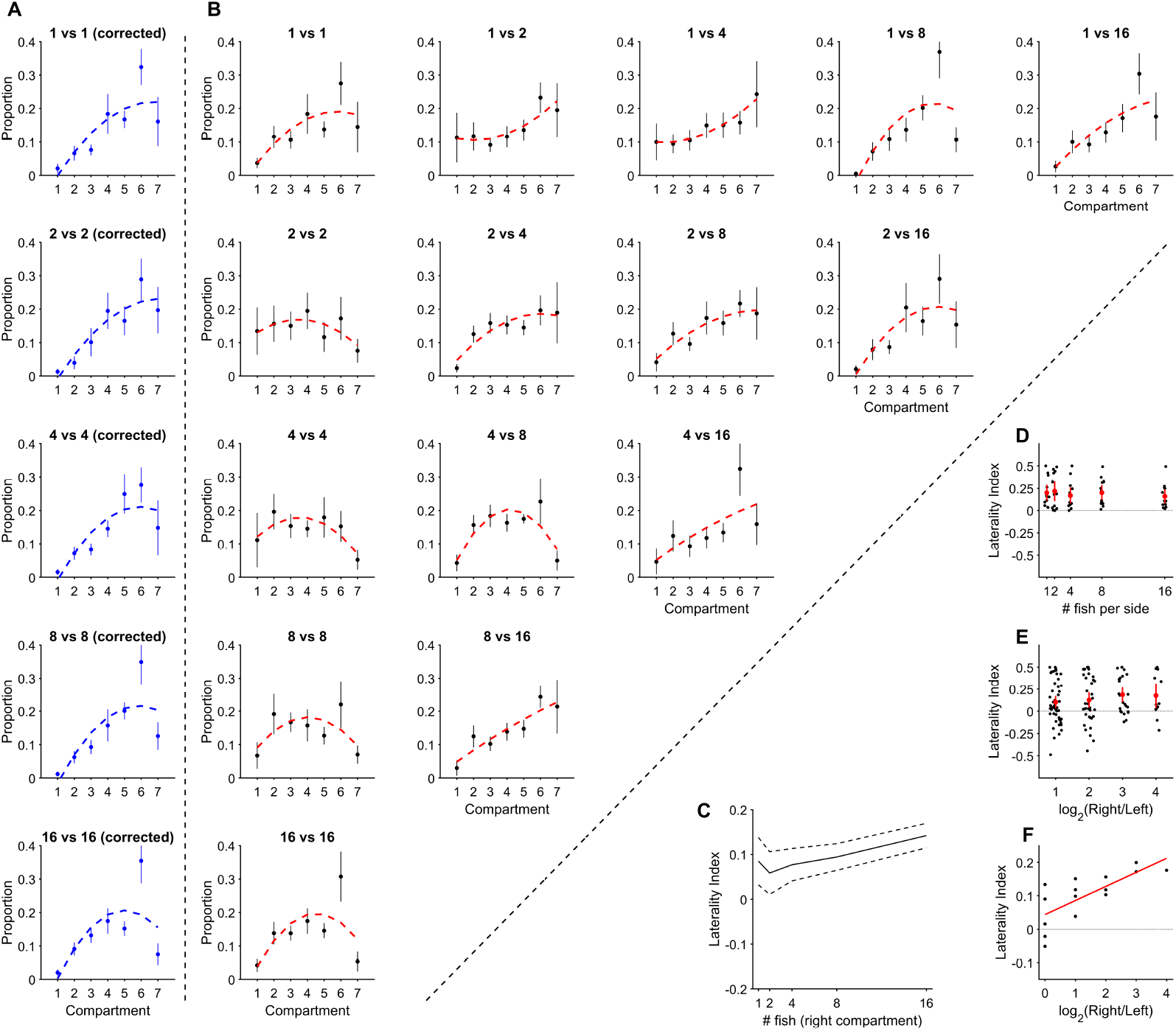
Compartment occupancy and laterality in two-choice tests with conspecific-like (small) stimuli. (A) Equal-number contrasts (e.g., 1 vs 1, 2 vs 2, …): profiles are realigned (trial-wise flipping) so that the chosen side is on the same end; error bars = SEM. (B) Raw mean compartment-occupancy profiles (SEM) for *both* the equal-number contrasts *without* choice alignment (e.g., 1 vs 1, 2 vs 2, …) and all non-equal contrasts (e.g., 1 vs 2, 1 vs 4, …) after magnitude alignment (larger numerosity assigned to Right). Red dashed curves are descriptive best-fitting polynomials. (C) Condition-level laterality index (LI) versus stimulus numerosity mapped to the comparison axis: for unequal-number contrasts this is the larger group’s numerosity; for equal-number contrasts it is the common numerosity. Curves show means with dashed mean±SEM envelopes. (D) Equal-number contrasts: choice-aligned LI (each trial mirrored so the chosen side is Right; descriptive), showing that fish typically chose one side rather than remaining central. Points with red mean ±95% CI. (E) Trial-level LI vs log2(R/L) for unequal-number contrasts (after magnitude alignment). Points with red mean ±95% CI for each bin. (F) Condition-mean LI vs log2(R/L); red line shows the population-level LME prediction (fixed effect).

**Figure 3:**
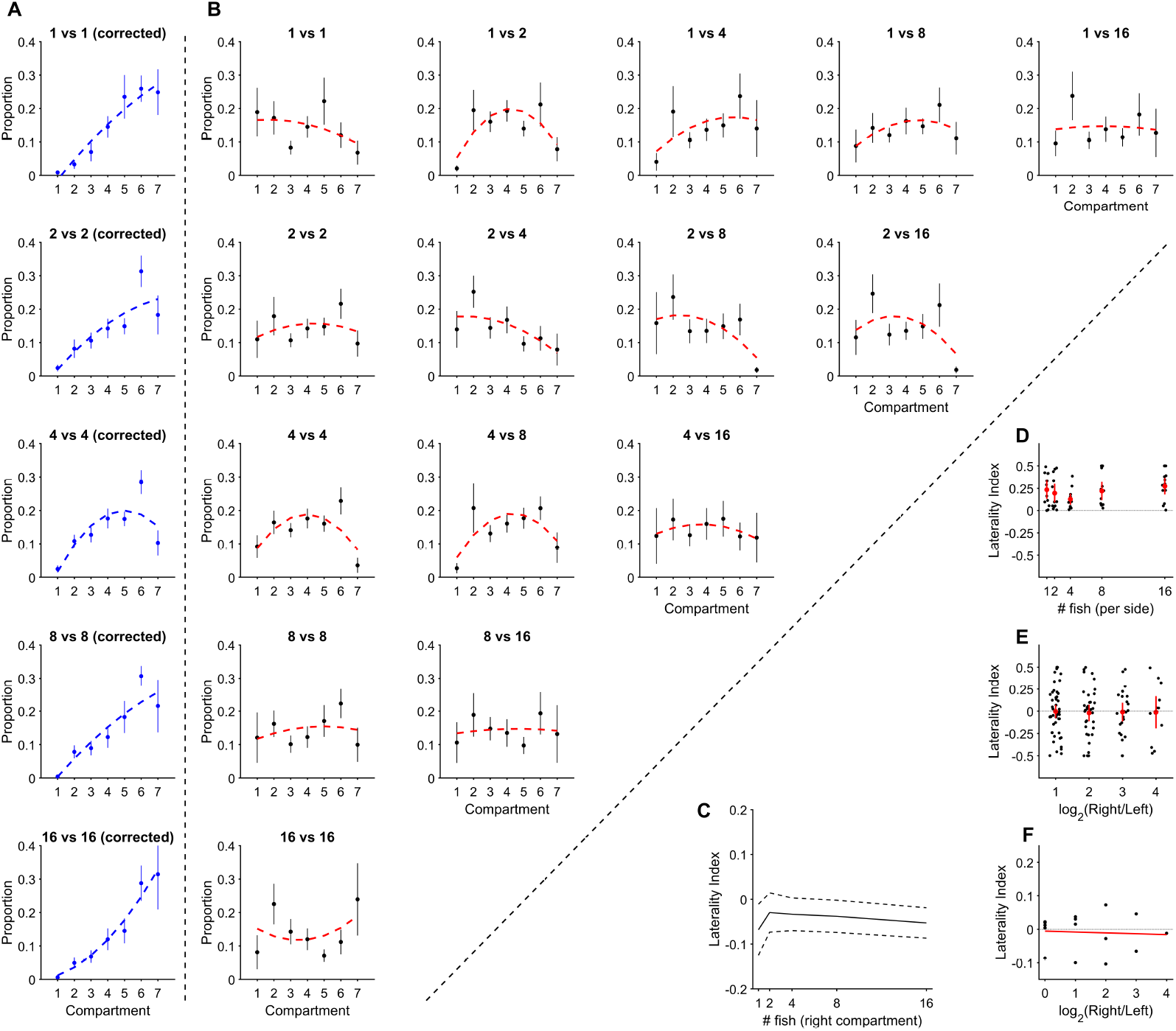
Compartment occupancy and laterality in two-choice tests with predator-like (large) stimuli. (A) Equal-number contrasts (e.g., 1 vs 1, 2 vs 2, …): profiles are realigned (trial-wise flipping) so that the chosen side is on the same end; error bars = SEM. (B) Raw mean compartment-occupancy profiles (SEM) for *both* the equal-number contrasts *without* choice alignment (e.g., 1 vs 1, 2 vs 2, …) and all non-equal contrasts (e.g., 1 vs 2, 1 vs 4, …) after magnitude alignment (larger numerosity assigned to Right). Red dashed curves are descriptive best-fitting polynomials. (C) Condition-level laterality index (LI) versus stimulus numerosity mapped to the comparison axis: for unequal-number contrasts this is the larger group’s numerosity; for equal-number contrasts it is the common numerosity. Curves show means with dashed mean±SEM envelopes. (D) Equal-number contrasts: choice-aligned LI (each trial mirrored so the chosen side is Right; descriptive), illustrating whether fish chose one side rather than remaining central. Points with red mean ±95% CI. (E) Trial-level LI vs log_2_(*R*/*L*) for unequal-number contrasts (after magnitude alignment). Points with red mean ±95% CI for each bin. (F) Condition-mean LI vs log_2_(*R*/*L*); red line shows the population-level LME prediction (fixed effect).

For *conspecific-like (small)* stimuli, laterality increased systematically with the numerical ratio: the fixed effect of log_2_(*N*_right_/*N*_left_) was positive and significant (*β* = 0.042 ± 0.013, *t*_178_ = 3.20, *p* = 0.0016; 95% CI [0.016, 0.068]), indicating an LI increase of ≈ 0.04 per doubling of the larger group relative to the smaller (Fig. 2E,F). The intercept did not differ from zero (*β*_0_ = 0.044 ± 0.038, *p* = 0.25). Fixed effects explained 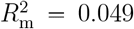 of the variance, rising to 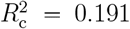 with random intercepts (12 subjects; *n* = 180 trials). The population fit increased monotonically across log_2_ ratios 0–4 (Fig. 2F). By contrast, for *predator-like (large)* stimuli the slope was near zero and non-significant (*β* = − 0.0026 ± 0.0162, *t* = − 0.16, *p* = 0.873; 95% CI [− 0.0345, 0.0293]), with both marginal and conditional *R*^2^ ≈ 0 (12 subjects; *n* = 180 trials), indicating no ratio-dependent laterality (Fig. 3E,F).

### Size × ratio interaction (trial-level LME)

Using the pooled interaction specification (Eq. 7) on two-choice trials with *N*_left_ > 0 and *N*_right_ > 0 (including equal-number contrasts, which contribute at log_2_(*N*_right_/*N*_left_) = 0), the analysis included *n* = 360 trials (small: 180, large: 180) from 12 subjects. Derived simple slopes showed clear ratio-dependence for conspecific-like stimuli (small: *β* = 0.0421, SE = 0.0153, 95% CI [0.0120, 0.0721]), but no ratio-dependence for predator-like stimuli (large: *β* = − 0.0026, SE = 0.0153, 95% CI [− 0.0326, 0.0274]). Critically, the size × log_2_ ratio interaction was negative (size_large_ × log_2_ ratio: *β* = − 0.0447, SE = 0.0213, *t*(356) = − 2.09, *p* = 0.037, 95% CI [−0.0866, −0.00270]), indicating that the ratio effect observed for conspecific-like stimuli was significantly reduced for predator-like stimuli. This conclusion was corroborated by a likelihood-ratio comparison against the corresponding main-effects model (*χ*^2^(1) = 4.35, *p* = 0.0369). As a secondary, non-independent check (trials aggregated to condition means; primary inference from the per-trial LME), we regressed condition-mean metrics on the log_2_ numerical ratio. For *small* stimuli, both the profile slope and the laterality index (LI) scaled with log_2_ ratio across the 15 contrasts: slope vs. log_2_ ratio, 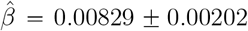, *t* = 4.10, *p* = 0.00125, adjusted *R*^2^ = 0.531; LI vs. log_2_ ratio, 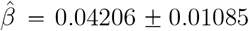, *t* = 3.88, *p* = 0.00190, adjusted *R*^2^ = 0.501. For *large* stimuli, neither slope nor LI showed a reliable relation to log_2_ ratio (slope 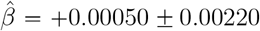, *p* = 0.824, adjusted *R*^2^ ≈ 0; LI 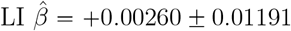, *p* = 0.831, adjusted *R*^2^ ≈ 0). Descriptive contrast-level parameters for each pairing are summarized in Table 3 (small stimuli) and Table 4 (large stimuli).

**Table 3:** Two-choice contrasts with conspecific-like (small) stimuli. Per-contrast mean LI, slope, curvature and *R*^2^, plus choice-aligned (^*^) fits for equal-number contrasts. *N* = 12 subjects.

| Contrast | $\log_2(R/L)$ | LI | *All contrasts*<br>*(unequal magnitude-aligned)* | | | *Equal-number only*<br>*(choice-aligned)* | | |
| --- | --- | --- | --- | --- | --- | --- | --- | --- |
| | | | Slope | Curv. | $R^2$ | Slope* | Curv.* | $R^{2*}$ |
| **1 vs 1** | 0.00 | 0.134 | 0.024 | -0.007 | 0.613 | 0.037 | -0.007 | 0.682 |
| **1 vs 2** | 1.00 | 0.099 | 0.019 | 0.005 | 0.735 | — | — | — |
| **1 vs 4** | 2.00 | 0.103 | 0.021 | 0.004 | 0.900 | — | — | — |
| **1 vs 8** | 3.00 | 0.199 | 0.035 | -0.011 | 0.556 | — | — | — |
| **1 vs 16** | 4.00 | 0.176 | 0.033 | -0.003 | 0.701 | — | — | — |
| **2 vs 2** | 0.00 | -0.021 | -0.006 | -0.006 | 0.490 | 0.040 | -0.006 | 0.841 |
| **2 vs 4** | 1.00 | 0.118 | 0.022 | -0.005 | 0.828 | — | — | — |
| **2 vs 8** | 2.00 | 0.118 | 0.024 | -0.004 | 0.828 | — | — | — |
| **2 vs 16** | 3.00 | 0.172 | 0.032 | -0.008 | 0.718 | — | — | — |
| **4 vs 4** | 0.00 | -0.051 | -0.008 | -0.009 | 0.658 | 0.035 | -0.009 | 0.753 |
| **4 vs 8** | 1.00 | 0.038 | 0.005 | -0.015 | 0.686 | — | — | — |
| **4 vs 16** | 2.00 | 0.156 | 0.028 | -0.001 | 0.475 | — | — | — |
| **8 vs 8** | 0.00 | 0.016 | 0.001 | -0.010 | 0.397 | 0.036 | -0.010 | 0.625 |
| **8 vs 16** | 1.00 | 0.151 | 0.030 | -0.001 | 0.834 | — | — | — |
| **16 vs 16** | 0.00 | 0.090 | 0.014 | -0.013 | 0.407 | 0.025 | -0.013 | 0.465 |

Curv. = quadratic coefficient; $R^2$ = quadratic fit goodness-of-fit. \* Choice-aligned descriptive fits are computed by mirroring equal-number trials to align the chosen side.

**Table 4:** Two-choice contrasts with predator-like (large) stimuli. Per-contrast mean LI, slope, curvature and *R*^2^, plus choice-aligned (^*^) fits for equal-number contrasts. *N* = 12 subjects.

| Contrast | $\log_2(R/L)$ | LI | *All contrasts*<br>*(unequal magnitude-aligned)* | | | *Equal-number only*<br>*(choice-aligned)* | | |
| --- | --- | --- | --- | --- | --- | --- | --- | --- |
| | | | Slope | Curv. | $R^2$ | Slope* | Curv.* | $R^{2*}$ |
| **1 vs 1** | 0.00 | -0.087 | -0.012 | -0.002 | 0.231 | 0.048 | -0.002 | 0.938 |
| **1 vs 2** | 1.00 | 0.037 | 0.007 | -0.014 | 0.603 | — | — | — |
| **1 vs 4** | 2.00 | 0.073 | 0.016 | -0.005 | 0.378 | — | — | — |
| **1 vs 8** | 3.00 | 0.046 | 0.008 | -0.005 | 0.468 | — | — | — |
| **1 vs 16** | 4.00 | -0.012 | 0.000 | -0.001 | 0.007 | — | — | — |
| **2 vs 2** | 0.00 | 0.012 | 0.003 | -0.004 | 0.114 | 0.035 | -0.004 | 0.713 |
| **2 vs 4** | 1.00 | -0.100 | -0.018 | -0.004 | 0.521 | — | — | — |
| **2 vs 8** | 2.00 | -0.104 | -0.019 | -0.006 | 0.532 | — | — | — |
| **2 vs 16** | 3.00 | -0.066 | -0.012 | -0.008 | 0.300 | — | — | — |
| **4 vs 4** | 0.00 | 0.004 | -0.001 | -0.012 | 0.481 | 0.023 | -0.012 | 0.652 |
| **4 vs 8** | 1.00 | 0.030 | 0.008 | -0.012 | 0.515 | — | — | — |
| **4 vs 16** | 2.00 | -0.028 | -0.002 | -0.004 | 0.377 | — | — | — |
| **8 vs 8** | 0.00 | 0.020 | 0.005 | -0.002 | 0.089 | 0.042 | -0.002 | 0.838 |
| **8 vs 16** | 1.00 | 0.015 | 0.001 | -0.001 | 0.016 | — | — | — |
| **16 vs 16** | 0.00 | 0.022 | 0.006 | 0.006 | 0.146 | 0.053 | 0.006 | 0.963 |

Curv. = quadratic coefficient; $R^2$ = quadratic fit goodness-of-fit. \* Choice-aligned descriptive fits are computed by mirroring equal-number trials to align the chosen side.

### Control analyses with image-based covariates

To test whether apparent numerical laterality could be explained by low-level stimulus properties, we re-analysed behaviour while adjusting for image-derived covariates (see Methods).

#### Trial-level GLMM (choice of the larger numerosity)

We fit a single binomial GLMM with logit link including *z*-scored log_2_ ratio (zLogRatio), *Size* (small/conspecific-like line drawings vs. large/predator-like line drawings), their interaction (zLogRatio×Size), and image covariates (ΔCoverage, ΣCoverage, and ΔNN with a missingness indicator), with random intercepts and random slopes for zLogRatio by subject (Eq. 8). Because numerosity and coverage were naturally correlated in our displays, this specification is deliberately conservative: zLogRatio competes with highly collinear coverage terms, trading off sensitivity to numerical effects against stringent control for simple continuous magnitudes. In the fitted model, none of the continuous coverage or spacing covariates emerged as strong predictors and neither the main effect of zLogRatio nor the zLogRatio×Size interaction reached significance (no fixed-effect *p* <.10), although their coefficients remained in the predicted directions and were attenuated relative to the LI analysis. Given the limited number of trials per contrast and the large set of covariates, this GLMM is underpowered to provide a precise estimate of the numerical effect, but it does indicate that simple cumulative area or global spacing statistics are unlikely to fully explain the pattern of choices. We therefore interpret it primarily as a stringent control analysis rather than as our most sensitive test of ratio-dependent preferences.

#### Condition-level regression (laterality index, LI)

For small-size stimuli, the image-based covariate model showed no reliable effects of log_2_(ratio) (*β* = − 0.0360 ± 0.0368, *p* = 0.351), ΔCoverage (*β* = − 0.405 ± 2.936, *p* = 0.893), ΣCoverage (*β* = 0.434 ± 0.770, *p* = 0.585), or Δ*NN* (*β* = − 1.78 × 10^-5^ ± 1.53 × 10^-4^, *p* = 0.910). For large-size stimuli, log_2_ (ratio) remained non-significant (*β* = 0.0475 ± 0.0342, *p* = 0.195), as did ΔCoverage (*β* = − 0.370 ± 0.471, *p* = 0.450) and ΣCoverage (*β* = 0.154±0.0988, *p* = 0.149), while Δ*NN* was negative and significant (*β* = − 3.44 × 10^-4^ ± 1.11× 10^-4^, *p* = 0.0115). In the large-stimulus condition, the local-spacing asymmetry covariate (ΔNN) was associated with laterality. However, controlling for the image-derived covariates did not remove the size-dependent ratio pattern.

### Occupancy profiles

To summarize spatial occupancy, we fitted linear and quadratic models to the mean compartment-occupancy profile for each contrast. For unequal-number contrasts (*N*_left_ ≠ *N*_right_), profiles were *magnitude-aligned* by swapping left/right labels as needed so that the larger numerosity is on the right before averaging. For equal-number contrasts (*N*_left_ = *N*_right_), no magnitude alignment is applicable. These contrasts are therefore reported as recorded. In addition, for equal-number contrasts we computed *choice-aligned* descriptive means by mirroring each trial so that the *chosen* side falls on the same side before averaging. This choice-aligned summary quantifies the strength of side commitment (approach a group vs. remain central) without implying a correct direction. Unless stated otherwise, tables report fits to *magnitude-aligned* means for unequal-number contrasts and to *unaligned* means for equal-number contrasts. *Choice-aligned* descriptive fits are reported separately for equal-number contrasts (Tables 3 and 4).

For conspecific-like (small) stimuli, unequal-number contrasts with larger disparities (e.g., 1 vs. 8, 1 vs. 16, 2 vs. 16) showed increasingly positive linear slopes, consistent with a shift toward the larger group, whereas curvature terms were small or negative, indicating largely monotonic profiles. For equal-number contrasts, raw means showed little consistent population-level side bias, whereas choice-aligned summaries yielded larger fitted slopes and *R*^2^ values, indicating that fish typically committed to one side even when both groups were identical.

### Ratio scaling and pooled tests (unequal-number contrasts)

At the pooled level (unequal-number contrasts only), laterality was reliably greater than zero for small stimuli (median LI = 0.049, *n* = 180, *p* < 0.0001), and both slope and LI increased with log_2_ ratio (contrast-level regressions: slope vs. log_2_ ratio, adjusted *R*^2^ = 0.546, *p* = 9.9 × 10^-4^; LI vs. log_2_ ratio, adjusted *R*^2^ = 0.511, *p* = 0.00164; Fig. 2E,F). For predator-like (large) stimuli, profiles showed no numerosity-driven affiliation: linear slopes clustered near zero across contrasts (range − 0.018 to 0.016; Table 4), pooled laterality did not differ from zero (median LI = − 0.006, *n* = 180, *p* = 0.676), and neither slope nor LI scaled with log_2_ ratio (contrast-level regressions: slope vs. log_2_ ratio, adjusted *R*^2^ ≈ 0, *p* = 0.864; LI vs. log_2_ ratio, adjusted *R*^2^ ≈ 0, *p* = 0.904; Fig. 3E,F).

### Model comparison: approximate number system vs object tracking system

We next asked whether trial-level preferences under unequal-number contrasts are better explained by Weber-like ratio scaling (ANS) or by an additional small-number sensitivity (OTS). We analysed unequal-number contrasts only (ratio > 1; equal-number trials excluded), and *z*-scored log_2_ ratios for numerical conditioning. For small stimuli, the ANS model (LI ∼ log_2_ ratio) had the lowest AIC (3.94), with ΔAIC relative to OTS = 1.44 and to the combined model = 1.97. Akaike weights were *w*_ANS_ = 0.54, *w*_OTS_ = 0.26, and *w*_ANS+OTS_ = 0.20, indicating modest preference for ANS. Adjusted *R*^2^ values were near zero (ANS = 0.009, OTS = − 0.003, combined = 0.000), consistent with the small effect sizes observed in the mixed-effects analyses. The smallSet term was not significant in the combined model (*p* = 0.864), arguing against an OTS-like sensitivity beyond ratio scaling in this regime. In contrast, for large stimuli neither predictor captured meaningful variance (ANS: AIC = 32.36, adj. *R*^2^= − 0.008; OTS: AIC = 32.23, adj. *R*^2^ = − 0.007; combined: AIC = 34.23, adj. *R*^2^ = − 0.016), and the smallSet term remained non-significant (*p* = 0.721), indicating that numerosity cues were not used to guide choice under predator-like stimuli.

## Discussion

In summary, zebrafish showed a clear, spontaneous preference to affiliate with the larger of two groups when the stimuli were conspecific-like. This preference scaled with the numerical ratio, consistent with a Weber-like approximate number system and providing no evidence for an additional small-number (OTS) advantage under our conditions. When the same numerosities were presented as predator-like stimulus images, numerosity ceased to guide two-choice behaviour. In single-option (0 vs. *N*) trials, laterality shifted reliably below zero, consistent with reduced approach to large-fish displays. This indicates that the numerical rule, i.e. the ANS-like decision heuristic, is flexibly gated by ecological context. Image-derived controls (coverage and nearest-neighbour spacing) did not provide a magnitude-only explanation for the pattern. Finally, in equal-number contrasts, choice-aligned averages showed that fish typically selected one group rather than remaining centrally. Predator-like stimulus appearance (large-fish displays; size with correlated morphology/identity differences) acts as a context-dependent switch: conspecific-like stimuli engage affiliation preference, whereas large-fish displays bias behaviour toward caution (most clearly in the 0 vs. *N* assay), thereby attenuating or reversing the usual preference for larger groups. That interpretation aligns with predation–risk frameworks and risk-sensitive group association, where stimulus meaning (e.g., apparent body size) modulates approach–avoidance decisions. Accordingly, we adopt a conservative interpretation: we treat the behavioural shift as a cautious response to unusually large, fish-like stimuli that likely signal elevated risk, rather than as direct evidence for predator recognition per se. This context-dependent gating aligns well with zebrafish work showing that purely visual predator cues can robustly alter behaviour: animated images of a sympatric predator and other fear-inducing stimuli elicit strong, stimulus-specific avoidance and freezing responses (Gerlai et al., 2009; Luca and Gerlai, 2012). At the group level, wild zebrafish adjust their group association decisions under predator risk, increasing association with mixed-species groups that may provide antipredator benefits (Mukherjee and Bhat, 2023). Our data extend these findings by showing that, under static visual stimulation in a spontaneous-choice task, the same numerical disparities that promote affiliation with conspecific-like groups no longer drive approach when apparent body size moves into a predator-like range. This interpretation aligns with frameworks showing that uncertainty about predation risk can bias behaviour toward caution and stronger defensive responding even when cues are indirect or ambiguous (Crane et al., 2024).

### Ratio-dependent quantity discrimination with conspecific-like stimuli

*H1 was supported:* with conspecific-like stimuli, zebrafish expressed a systematic, monotonic preference for the side depicting the larger group. The strength of this preference scaled with the log_2_ ratio between numerosities. This pattern is the hallmark of an approximate number system (ANS): equal ratios (e.g., 1 vs. 2) elicit comparable preference strength across absolute set sizes, consistent with Weber-like coding of relative, rather than absolute, differences. In our model comparisons, a ratio term accounted for the systematic component of laterality, whereas adding a small-number factor (coding pairs drawn only from {1,2,4}) did not provide additional explanatory value. Thus, within the range sampled in this study, behaviour aligned with ANS-like ratio dependence without evidence for an extra advantage confined to very small sets.

These findings align with spontaneous-choice results in fish showing that preferences for larger groups typically follow ratio limits (e.g., Agrillo et al., 2010; Gómez-Laplaza and Gerlai, 2011; Agrillo et al., 2012; Agrillo and Bisazza, 2018). At the same time, reports of disproportionate performance at very small numerosities (“subitizing”-like advantages, i.e. a fast, accurate identification of small numerosities (≈ 1–4) without serial counting) do exist, particularly in designs that tightly equalise non-numerical cues such as cumulative area, convex hull, spacing, and perimeter (Gebuis and Reynvoet, 2011; Agrillo and Bisazza, 2018). Our stimuli were ecologically styled and not cue-matched in that strict sense. Accordingly, the lack of detectable small-number sensitivity here should not be taken as evidence against OTS. Rather, it suggests that under naturalistic viewing with heterogeneous visual structure and after adjusting for major image covariates zebrafish behave as if guided primarily by relative numerosity.

Finally, the ANS-consistent scaling in the conspecific-like condition also clarifies the boundary we observe when the stimulus regime switches to predator-like large-fish line drawings. The same ratio structure that supports affiliation with small fish stimuli fails to drive approach with predator-like fish stimuli (see below), indicating that ANS-like evaluation is not a fixed default but is expressed when the ecological context favours social affiliation. This situates zebrafish alongside other taxa in which ratio dependence is reliable yet gated by motivational state and stimulus meaning, rather than being an invariant output of low-level vision (Feigenson et al., 2004; Cantlon and Brannon, 2007; Beran, 2007; Rugani et al., 2015).

### Apparent body size modulates the use of numerosity

*H2 was supported:* When the same numerical contrasts were presented using large-fish (predator-like) line drawings, the preference for the larger group that characterised conspecific-like stimuli disappeared. Moreover, in the single-option (0 vs. *N*) assay laterality shifted reliably below zero, consistent with a cautious response to large-fish displays. This pattern is difficult to reconcile with a purely low-level magnitude account: although the two stimulus sets differed in fish-model morphology/identity (and corresponding apparent body size), they shared the same numerical manipulation, and continuous quantities that co-vary with number, including the image-derived features we quantified (coverage and nearest-neighbour spacing), varied systematically with numerosity in *both* stimulus sets. Yet only the conspecific-like displays elicited numerosity-guided approach. A parsimonious interpretation is therefore that predator-like stimulus appearance shifts the *decision context* from affiliation to risk management: with conspecific-like displays, zebrafish treat the drawings as social opportunities and favour approach toward the numerically larger group; with large-fish displays, the same quantitative regularities (numerical contrast and its correlated magnitudes) are down-weighted and behaviour is biased toward caution, attenuating (or abolishing) numerosity-based approach. This account accords with predation-risk theory and risk-sensitive group association (Lima and Dill, 1990; Kelley and Magurran, 2003; Krause and Ruxton, 2002).

At the circuit level, apparent body size and predator-like morphology likely shift the balance between social-approach and threat-avoidance systems. Conspecific-like line drawings may bias processing toward affiliative networks, including preoptic and hypothalamic social circuits and dopaminergic valuation pathways, allowing numerosity to guide approach. Predator-like large-fish line drawings may instead engage size- and threat-sensitive visual pathways in the pretectum and tectum (Temizer et al., 2015; Dunn et al., 2016), recruiting downstream avoidance and arousal systems such as reticulospinal neurons and stress-related modulators that suppress approach behaviour. This interpretation is consistent with established zebrafish visual–motor circuitry and predation-risk frameworks (Lima and Dill, 1990), explaining why the same numerical structure promotes affiliation in the conspecific-like condition but not in the large-fish condition. Large angular size (and, more generally, stimulus features characteristic of potential threats) can engage escape and vigilance circuitry, biasing behaviour away from nearby, visually salient masses (Dunn et al., 2016; Fotowat and Engert, 2023). In zebrafish, number-sensitive changes in gene expression have been found in the telencephalon and thalamus when the number of visual items changes. However, changes in overall stimulus size evoke stronger responses in the optic tectum/retina, supporting the idea of partially distinct processing pathways for numerosity vs. magnitude cues (Messina et al., 2020).

Conceptually, these results suggest a context-dependent gating mechanism: numerosity is not an invariant driver of approach but one cue among many whose influence is modulated by decision context and stimulus meaning. This yields testable predictions. First, reducing perceived risk (e.g., providing a refuge or a familiar background) should attenuate the negative laterality/reduced approach to large-fish displays; whether this is sufficient for ratio-based affiliation to reappear remains an open empirical question. Second, other threat cues (e.g., eyespots, contrast polarity, looming motion) should similarly suppress numerosity-guided approach even when continuous magnitudes favour the larger side. Together, the data argue for a context-sensitive integration of quantitative and ethological information rather than a fixed, low-level magnitude heuristic.

### Equal-number trials indicate active affiliation

*H3 was supported for conspecific-like stimuli:* When the two sides displayed the same numerosity in the conspecific-like condition, raw group means showed little directional bias at the population level, as expected, because opposite choices cancel in the average. Choice-aligned profiles address this averaging artifact by mirroring individual trials so that the *chosen* side is plotted consistently. The resulting distributions peak near the chosen side rather than at the tank centre, indicating that fish typically made an active affiliation decision even without a numerical advantage. This rules out a trivial “no decision/loitering” account for equal-number trials with conspecific-like stimuli and confirms that the assay elicits genuine approach–avoid behaviour. In contrast, for predator-like displays, equal-number trials more often showed central occupancy and weak or absent side selection, reinforcing the interpretation that apparent size shifts the decision context away from affiliation and toward caution.

### Implications for numerosity-guided decision making

H4 was broadly consistent with the data: With conspecific-like stimuli, preference strength increased with the logarithmic ratio between groups, and this Weber-like pattern was well captured by an ANS-like model. Adding a small-number (OTS-like) term did not improve explanatory power in this spontaneous-choice context. However, our stimuli were not specifically engineered to orthogonalise numerosity from continuous magnitudes (e.g., cumulative area, density, convex hull) in the 1–4 range. Under these conditions, we therefore find no detectable additional benefit confined to very small numerosities. But we cannot exclude the possibility of a distinct small-number mechanism in zebrafish. Two features of the design may contribute: First, choices were untrained and embedded in a social setting, where approximate, ratio-based valuation is likely sufficient and efficient. Second, ecological context modulates whether number guides action at all, i.e., stimuli consisting of predator-like large-fish line drawings reduce approach to the numerically larger group. Any subtle small-number advantage could thus be masked by such context-dependent policies. Because trials were distributed across a broad range of ratios and both stimulus regimes, rather than concentrated on a few very small-number contrasts, some fine-grained effects (e.g. putative OTS-specific advantages restricted to 1–4, or subtle biases on equal-numerosity trials by stimulus regime) are only moderately powered. Null or weak effects in these sub-analyses should therefore be taken as an absence of clear evidence rather than definitive evidence of absence. A conservative reading is that zebrafish show an ANS-like, ratio-dependent behavioural signature in spontaneous affiliation choices in our task, with no evidence of an additional OTS-like advantage under these conditions. If such an advantage exists, it may only emerge under tighter cue control, greater within-subject power, or tasks that explicitly require exact judgments of small sets.

Our data are consistent with an approximate, ratio-sensitive representation of conspecific group size in zebrafish, modulated by the ecological meaning of the stimulus. At the same time, work in other teleosts suggests that such numerical representations can be implemented even in highly compact brains: in the miniature, transparent *Danionella cerebrum*, Zanon et al. (2025) demonstrated quantity discrimination for dot arrays under stringent control of basic continuous magnitudes. Together, these findings support the idea that basic numerical competence is a widespread capacity in teleosts, while our results add that the influence of number on behaviour is flexibly gated by context, such as predator-like stimulus appearance (including apparent body size and morphology) and putative predation risk.

### Low-level magnitudes do not account for behaviour across stimulus regimes

We asked whether simple image magnitudes could account for the behavioural patterns by adding coverage and spacing covariates (Eq. 1) to the models (Eqs. 8, 9). For *conspecific-like* stimuli, neither the coverage *difference* (ΔCoverage) nor the *sum* (ΣCoverage), and likewise not nearest-neighbour spacing, predicted laterality or choice once numerical ratio was included. Thus, larger-group affiliation in this regime is not reducible to cumulative area or simple density. For *large-fish (predator-like)* stimuli, coverage terms again carried little explanatory value: only a small, negative ΔNN coefficient emerged, indicating that tighter spacing on the side with more (large) items was associated with a slightly lower laterality index. In other words, denser clustering of large-fish figures modestly biased fish away from that side. Crucially, this spacing effect did not reinstate numerosity use: ratio did not guide choice in the large-fish regime. Together, these controls show that simple coverage-based magnitudes cannot explain the behavioural dissociation between stimulus regimes, and while spacing may modulate caution/avoidance under predator-like stimulus appearance (large-fish line drawings; morphology/identity and size), it does not recover numerosity-guided approach behaviour.

## Limitations of the study

Two limitations qualify the interpretation of the stimulus-regime effect. First, we did not include an independent manipulation check demonstrating that the large-fish displays are categorised as predators (or otherwise explicitly threatening) by the animals. Accordingly, we interpret the large-fish condition conservatively as a predator-like/threat-like visual regime rather than as evidence of predator recognition per se. Second, the large-fish regime differs from the small-fish regime not only in apparent size but also in correlated stimulus properties such as morphology/identity, meaning that we did not isolate size from other regime-defining visual features. Throughout, references to “size” should therefore be read as shorthand for this predator-like regime. Finally, our stimuli were static 2D line drawings, which may not capture key cues present in natural encounters (e.g., motion, looming, or 3D structure), and future work should test whether the same context-dependent gating of numerosity-guided affiliation generalises to more naturalistic and independently validated threat manipulations.

## Conclusion

Zebrafish exhibit spontaneous, ratio-dependent quantity discrimination when stimuli are conspecific-like (small), but the same numerical cues no longer govern choices when the stimuli are predator-like (large), where numerosity no longer promotes approach. We did not isolate size from morphology; therefore ‘size’ is shorthand for the predator-like regime. In the single-option assay, large-fish displays elicited negative laterality consistent with a cautious response. Image-derived controls (coverage and nearest-neighbour spacing) did not account for these effects: simple differences in cumulative area or mean spacing were insufficient to explain behaviour across size regimes, and numerosity remained a key predictor in the conspecific-like regime. We cannot, of course, exclude contributions from more complex low-level statistics (e.g. fine-grained clustering), but within our stimulus set no single coverage or spacing metric could plausibly replace numerosity as the main driver of behaviour. Consistent with this, a deliberately conservative trial-level GLMM that jointly included logarithmic ratio and image-derived coverage and spacing covariates did not identify any simple continuous metric that could replace numerosity as the main driver of choices. Our data therefore argue against a simple magnitude-only account based on cumulative area or density alone, and instead support a view in which numerosity is one cue whose influence is modulated by the ecological meaning of the display. Together, the findings suggest that numerosity acts as one cue in a broader decision process whose influence is modulated by ecological meaning. That is, in social contexts it supports approach toward larger groups, whereas in threat-like contexts it is suppressed in favour of caution. This flexible weighting offers a principled bridge between quantitative decision rules and ethologically appropriate action, and motivates future work to map the size boundary of this modulation and test its neural implementation.

## Acknowledgements

We are grateful for the financial support by the Startup-funding of Taipei Medical University (TMU108-AE1-B33) and the Taiwan Ministry of Science and Technology research grant (110-2311-B-038-002, 112-2410-H-038-027) granted to CDD. We thank Jiun-Lin Horng for providing access to the experimental facilities that made this work possible and for technical support during data collection.

## Data and code availability

All analysis code used in this study is publicly available at https://github.com/ChristophDahl/fishQuantity/.

The data supporting the findings of this study, including behavioural datasets and derived stimulus metrics, are available on the Open Science Framework (OSF) at https://osf.io/wndsm/overview/.

## Author contributions

HC: conceptualisation, study design, data collection, writing.

CDD: conceptualisation, study design, data collection, analysis and interpretation, provision of necessary tools/software, writing.

Both authors approved the final version of the manuscript.

## Ethical approval

The experimental protocol was reviewed and approved by the Animal Care and Use Committees of Taipei Medical University (LAC-2021-0660, LAC-2025-0214).

## References

Addessi E, Crescimbene L, Visalberghi E (2008) Food and token quantity discrimination in capuchin monkeys (Cebus apella). Animal Cognition 11(2):275–282

Agrillo C, Bisazza A (2018) Understanding the origin of number sense: a review of fish studies. Philosophical Transactions of the Royal Society B: Biological Sciences 373(1740):20160511

Agrillo C, Dadda M, Serena G, Bisazza A (2008) Do fish count? spontaneous discrimination of quantity in female mosquitofish. Animal Cognition 11(3):495–503

Agrillo C, Piffer L, Bisazza A (2010) Large number discrimination by mosquitofish. PLoS One 5(12):e15232

Agrillo C, Piffer L, Bisazza A, Butterworth B (2012) Evidence for two numerical systems that are similar in humans and guppies. PLoS One 7(2):e31923

Agrillo C, Petrazzini MEM, Bisazza A (2017) Numerical abilities in fish: a methodological review. Behavioural Processes 141:161–171

Aslanzadeh M, Ariyasiri K, Kim OH, Choi TI, Lim JH, Kim HG, Gerlai R, Kim CH (2019) The body size of stimulus conspecifics affects social preference in a binary choice task in wild-type, but not in dyrk1aa mutant, zebrafish. Zebrafish 16(3):262–267

Beran MJ (2007) Rhesus monkeys (Macaca mulatta) enumerate large and small sequentially presented sets of items using analog numerical representations. Journal of Experimental Psychology: Animal Behavior Processes 33(1):42

Bisazza A, Piffer L, Serena G, Agrillo C (2010) Ontogeny of numerical abilities in fish. PLoS One 5(11):e15516

Cantlon JF, Brannon EM (2007) How much does number matter to a monkey (Macaca mulatta)? Journal of Experimental Psychology: Animal Behavior Processes 33(1):32

Crane AL, Feyten LE, Preagola AA, Ferrari MC, Brown GE (2024) Uncertainty about predation risk: a conceptual review. Biological Reviews 99(1):238–252

Dadda M, Piffer L, Agrillo C, Bisazza A (2009) Spontaneous number representation in mosquitofish. Cognition 112(2):343–348

Dehaene S (2011) The number sense: How the mind creates mathematics. OUP USA

Dunn TW, Gebhardt C, Naumann EA, Riegler C, Ahrens MB, Engert F, Del Bene F (2016) Neural circuits underlying visually evoked escapes in larval zebrafish. Neuron 89(3):613–628

Feigenson L, Dehaene S, Spelke E (2004) Core systems of number. Trends in Cognitive Sciences 8(7):307–314

Fernandes Y, Rampersad M, Jia J, Gerlai R (2015) The effect of the number and size of animated conspecific images on shoaling responses of zebrafish. Pharmacology Biochemistry and Behavior 139:94–102

Ferrando E, Dahl CD (2022) An investigation on the olfactory capabilities of domestic dogs (Canis lupus familiaris). Animal Cognition 25(6):1567–1577

Fotowat H, Engert F (2023) Neural circuits underlying habituation of visually evoked escape behaviors in larval zebrafish. Elife 12:e82916

Gallistel CR, Gelman R (2000) Non-verbal numerical cognition: From reals to integers. Trends in Cognitive Sciences 4(2):59–65

Gebuis T, Reynvoet B (2011) Generating nonsymbolic number stimuli. Behavior Research Methods 43(4):981–986

Gebuis T, Reynvoet B (2012) The interplay between nonsymbolic number and its continuous visual properties. Journal of Experimental Psychology: General 141(4):642

Gerlai R, Fernandes Y, Pereira T (2009) Zebrafish (Danio rerio) responds to the animated image of a predator: towards the development of an automated aversive task. Behavioural Brain Research 201(2):318–324

Gómez-Laplaza LM, Gerlai R (2011) Can angelfish (Pterophyllum scalare) count? discrimination between different shoal sizes follows weber’s law. Animal Cognition 14(1):1–9

Horowitz A, Hecht J, Dedrick A (2013) Smelling more or less: Investigating the olfactory experience of the domestic dog. Learning and Motivation 44(4):207–217

Howard SR, Schramme J, Garcia JE, Ng L, Avarguès-Weber A, Greentree AD, Dyer AG (2020) Spontaneous quantity discrimination of artificial flowers by foraging honeybees. Journal of Experimental Biology 223(9):jeb223610

Jones SM, Brannon EM (2012) Prosimian primates show ratio dependence in spontaneous quantity discriminations. Frontiers in Psychology 3:550

Kelley JL, Magurran AE (2003) Learned predator recognition and antipredator responses in fishes. Fish and Fisheries 4(3):216–226

Krause J, Ruxton GD (2002) Living in Groups. Oxford Series in Ecology and Evolution, Oxford University Press, Oxford

Landeau L, Terborgh J (1986) Oddity and the ‘confusion effect’in predation. Animal Behaviour 34(5):1372–1380

Lima SL, Dill LM (1990) Behavioral decisions made under the risk of predation: a review and prospectus. Canadian Journal of Zoology 68(4):619–640

Luca RM, Gerlai R (2012) In search of optimal fear inducing stimuli: differential behavioral responses to computer animated images in zebrafish. Behavioural Brain Research 226(1):66–76

Messina A, Potrich D, Schiona I, Sovrano VA, Fraser SE, Brennan CH, Vallortigara G (2020) Response to change in the number of visual stimuli in zebrafish: A behavioural and molecular study. Scientific Reports 10(1):5769

Messina A, Potrich D, Schiona I, Sovrano VA, Vallortigara G (2021) The sense of number in fish, with particular reference to its neurobiological bases. Animals 11(11):3072

Mukherjee I, Bhat A (2023) What drives mixed-species shoaling among wild zebrafish? the roles of predators, food access, abundance of conspecifics and familiarity. Biology Open 12(1):bio059529

Nakagawa S, Schielzeth H (2013) A general and simple method for obtaining r2 from generalized linear mixed-effects models. Methods in Ecology and Evolution 4(2):133–142

Nieder A (2025) The calculating brain. Physiological Reviews 105(1):267–314

Nys J, Content A (2012) Judgement of discrete and continuous quantity in adults: Number counts! Quarterly Journal of Experimental Psychology 65(4):675–690

Ogi A, Licitra R, Naef V, Marchese M, Fronte B, Gazzano A, Santorelli FM (2021) Social preference tests in zebrafish: a systematic review. Frontiers in Veterinary Science 7:590057

Petrazzini MEM, Wynne CD (2016) What counts for dogs (Canis lupus familiaris) in a quantity discrimination task? Behavioural Processes 122:90–97

Peuhkuri N (1997) Size-assortative shoaling in fish: the effect of oddity on foraging behaviour. Animal Behaviour 54(2):271–278

Piffer L, Agrillo C, Hyde DC (2012) Small and large number discrimination in guppies. Animal Cognition 15(2):215–221

Pita D, Fernández-Juricic E (2021) Zebrafish neighbor distance changes relative to conspecific size, position in the water column, and the horizon: A video-playback experiment. Frontiers in Ecology and Evolution 8:568752

Potrich D, Montel L, Stancher G, Baratti G, Vallortigara G, Sovrano VA (2024) Proto-arithmetic abilities in zebrafish (Danio rerio). Heliyon 10(23)

Pritchard VL, Lawrence J, Butlin RK, Krause J (2001) Shoal choice in zebrafish, Danio rerio: the influence of shoal size and activity. Animal Behaviour 62(6):1085–1088

Qin M, Wong A, Seguin D, Gerlai R (2014) Induction of social behavior in zebrafish: live versus computer animated fish as stimuli. Zebrafish 11(3):185–197

Rugani R, Vallortigara G, Priftis K, Regolin L (2015) Number-space mapping in the newborn chick resembles humans’ mental number line. Science 347(6221):534–536

Saverino C, Gerlai R (2008) The social zebrafish: behavioral responses to conspecific, heterospecific, and computer animated fish. Behavioural Brain Research 191(1):77–87

Swaney WT, Jose A, Hirons-Major C, Reddon AR (2025) Decision-making in shoaling: zebrafish integrate cues of familiarity and group size. Animal Cognition 28(1):92

Temizer I, Donovan JC, Baier H, Semmelhack JL (2015) A visual pathway for looming-evoked escape in larval zebrafish. Current Biology 25(14):1823–1834

Ward C, Smuts BB (2007) Quantity-based judgments in the domestic dog (Canis lupus familiaris). Animal Cognition 10(1):71–80

Zanon M, Fraser SE, Vallortigara G (2025) Numerical discrimination in danionella. iScience 28(11)

